# Self-organization of cortical areas in the development and evolution of neocortex: a network growth model

**DOI:** 10.1101/2020.05.13.094672

**Authors:** Nabil Imam, Barbara Finlay

## Abstract

While the mechanisms generating the topographic organization of primary sensory areas in the neocortex are well-studied, what generates secondary cortical areas is virtually unknown. Using physical parameters representing primary and secondary visual areas as they vary from monkey to mouse, we derived a growth model to explore if characteristic features of secondary areas could be produced from correlated activity patterns arising from V1 alone. We found that V1 seeded variable numbers of secondary areas based on activity-driven wiring and wiring density limits within the cortical surface. These secondary areas exhibited the typical mirror-reversal of map topography on cortical area boundaries and progressive reduction of the area and spatial resolution of each new map on the caudorostral axis. Activity-based map formation may be the basic mechanism that establishes the matrix of topographically-organized cortical areas available for later computational specialization.

## 1 Introduction

In 1909, Brodmann divided the entire expanse of the human cerebral cortex into 52 “areas”, an analysis which organized research on the cortex for the following century [1]. Cortical areas were numbered in the order Brodmann encountered each new type in horizontal sections from the top to the bottom of the cortex. The histological evidence available to him included the presence and quantity of neurons and fiber layers, details of staining, characterization of cell body types and process elaborations of neurons in each area, and the numbers of non-neuronal cells, together called “cytoarchitectonics”.

Subsequent work, in description of connectivity and topographic representation [2–4], pharmacological and immunohistochemical characterization of neuronal types [5], electrophysiological characterization of single neuron properties, neuroimaging [6] and gene expression [7] largely have reified Brodmann’s divisions, though subdividing and elaborating his choices. Cortical areas became to be defined by a conjunction of interrelated properties. Each cytoarchitectonic area of cortex differs from its immediate neighbors in the particular thalamic nuclei, subcortical regions or intracortical areas it connects with. Each defined area could be further associated with a particular collection of electrophysiologically-defined receptive field types, ranging over Hubel and Wiesel’s edge detectors in primary visual cortex, “Area 17” [8], to a hierarchy of abstract decision properties in frontal cortex [9]. Central to the present study, cortical areas typically presented topologically-organized representations of sensory or motor surfaces like the retina or cochlea, secondary computed representations like intermodal egocentric space, or computed dimensions like “decision abstraction” [10]. By this confluence of dimensions, cortical areas retained the status of the central unit of cortical organization. By analogy to the electronics of the research era, each area was usually imagined as an input-output device that performed a particular transformation in accord with its unique within-area circuitry, passing on its results to other areas to be integrated with other inputs in a rough hierarchy first described by Van Essen and colleagues [11], often named according to their apparently dominant function.

Any typology defined by a loose aggregation of properties generates controversies. From the start, the uncertain relationship of neurological symptoms to the proposed function of areas (for example, the language and other functions of “Broca’s area” [12]), caused controversy on the computational centrality of the cortical area. Adjacent areas might have only unimpressive differences in the ratios of electrophysiological classes of neurons, contrasted with the distinct functional names assigned to them (e.g. “Color” vs “Motion” [13]). Neuroimaging expands the controversy, where varying methods of analysis can alternately distinguish unique functions associated with each area (e.g “Fusiform Face Area, FFA” [14]) or a near unlimited depth of distinct sensory, motor, or integrative functions reaching across specific areas (reviewed in [15, 16]). Influential network analyses, which typically define cortical areas as network “nodes”, can demonstrate new functional groupings over the classical typologies [17] but if metric distance as well as node “identity” linked to area is considered, different organizational principles emerge [18,19]. Comparing cortical organization in different species, where larger cortices usually present more and more “areas”, the question arises whether the new areas are add-ons, duplications, subdivisions, or complete reorganizations of larger defined zones [20–22].

A distinct developmental duality in the mechanisms by which cortical areas are positioned in the cortical surface and innervated has the potential to point at what mechanisms might generate non-primary cortical areas. Primary sensory and motor areas are distinct from all other areas by being recognizable at the earliest developmental stages, genomically, neuroanatomically and physiologically [7]. Primary sensory areas uniquely attract and recognize, trophically require and topographically organize input from their respective primary sensory thalamic nuclei with extreme specificity [21, 23–25] earlier than secondary cortical areas receive thalamic input [26]. These primary cortical areas, positioned on the overall cortical surface by diffusible gradients emanating from the rostral and caudal poles of the cortical plate [28], are often said to “organize” the cortical map. A curious absence in cortical research becomes evident at this point. Though literally thousands of studies have been performed on the development of the topology, connectivity and single unit response properties of primary sensory and motor areas (for example, [27] [28]), few to no such studies of “secondary” or “association” areas, particularly at the earliest stages of development now well-known for primary visual or somatosensory cortex have been done.

Decades of work characterizing the topography of the visual field representations in the occipital and parietal cortex [6, 29], coupled with a similar depth of work uncovering the multiple mechanisms of topographic map formation in the brain [30, 31] offer a way in to understand how non-primary cortical fields might develop (Figure 1A; redrawn from [32]). Primary visual cortex, V1, is the largest in surface area of the visual representations, and has a point-to-point representation of the retinal surface at high resolution. Secondary visual cortex, smaller in area, is topographically continuous with V1, mirror-reversing the V1 center-to-periphery retinotopic map at its anterior border while retaining its up-down polarity. “V3” is narrower still, again reversing polarity; further maps begin to fractionate. Overall, the anatomical and physiological topographic “resolution” of these maps decreases with distance from V1 [33].

**Figure 1:**
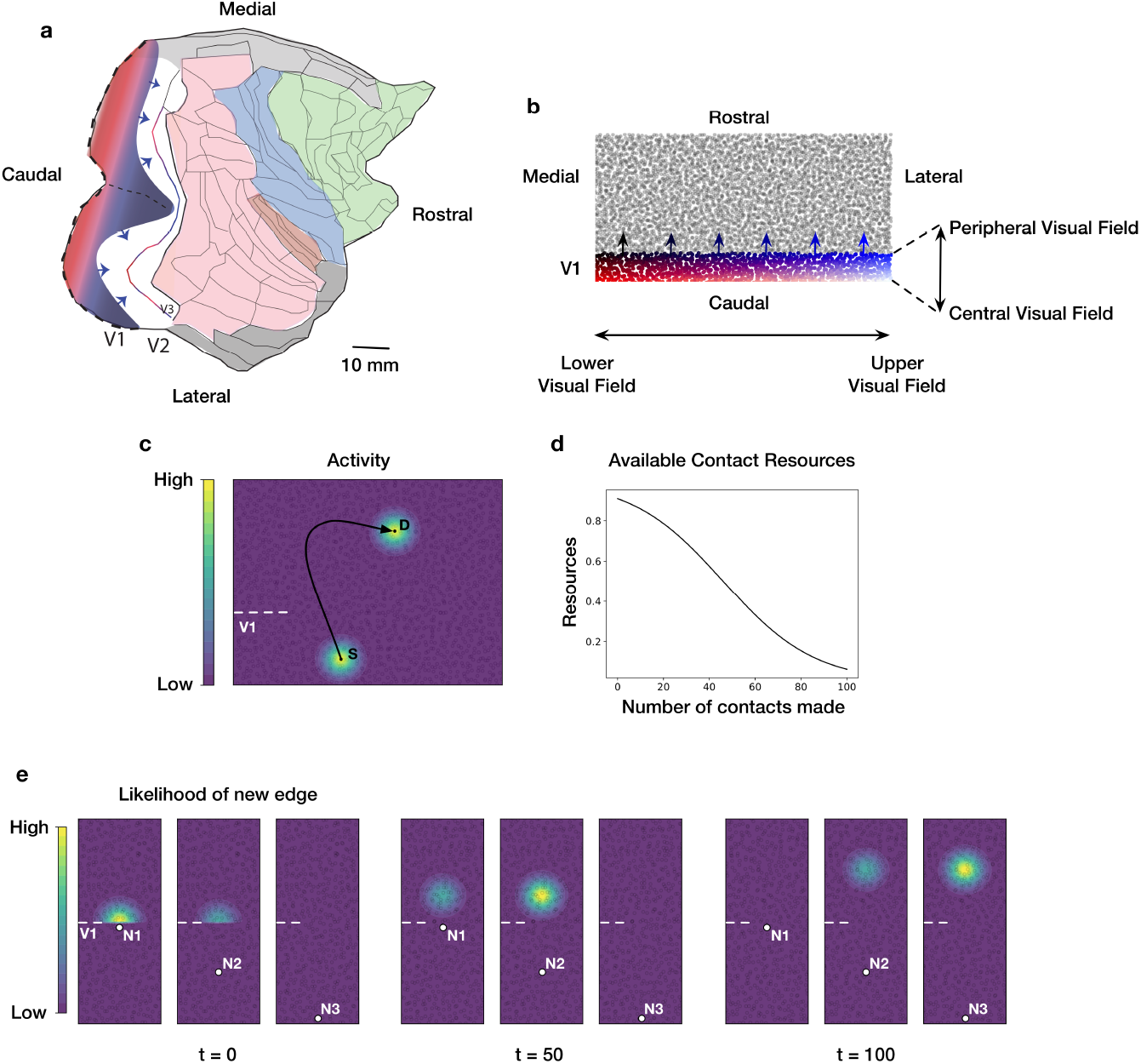
(a) Flattened representation of the right cortical hemisphere of a macaque monkey. Dotted line is where V1 is cut down the horizontal meridian to flatten it. Arrows show advancing axons exiting V1 into V2. The colored regions are frontal (green), limbic (grey), somatosensory (blue), and extendend occipital-interparietal cortex (pink). (b) The initial state of the network model. Individual nodes represent neural population under a 1 mm by 1 mm piece of cortex. The network grows in sequential steps by forming new directed edges from nodes in V1 to nodes in the rest of the cortical sheet. (c) Contour plot illustrating spread of neural activity in a representative piece of the model. Activity of V1 node S generates correlated activity in its immediate spatial neighborhood. This activity falls off as a Gaussian outwards from S. Activity in S also generates correlated activity on a distant node D to which S projects an edge to. Activity of D spreads to its neighborhood and falls off as a Gaussian outwards from D. At each growth step, new edges preferentially form between nodes that have high activity correlations. (d) As new edges are added to a node, its available contact (synaptic) resources decreases as a sigmoidal function. New edges preferentially form between nodes that have a larger number of available contact resources. (e) Contour plot in a representative piece of the model, illustrating the likelihood (probability) of a new edge emanating from one of three nodes (S1, S2, S3) in V1 and terminating in any of the nodes outside of V1. The distribution of probabilities are shown at three different time points. In the network’s initial state (time t=0, when no edges have yet formed), new edges are more likely to form between node S1 and nodes outside of V1 that are in immediate spatial proximity to S1. Subsequently at time t=50 (followed by t=100), new edges are more likely to form between node S2 (followed by S3) and nodes outside of V1 further along the caudorostral axis.

Secondary cortical areas could be generated directly by the early, topographically-organized and active axon innervation from primary cortical areas, using activity-driven mechanisms so amply demonstrated in the organization of binocular receptive fields, orientation selectivity and so forth in primary visual cortex, but largely missing the molecular axon/substrate recognition systems critical for the early emergence of V1 topography. Here we report on a network model of the visual cortex that self organizes based on activity-dependent correlations emanating from a single topographically-specified zone. We show that topographical properties of secondary visual areas, including mirror-symmetry and progressive change in map size and resolution arise from two features of the developing cortex: activity-based neuronal wiring and wiring density limits. We investigate variations of map features with changes in parameters specifying these factors, and analyze systematic changes in map organization with variations in overall brain size.

## 2 Growth Model

We used the relative dimensions of primary and secondary visual cortical regions of the rhesus macaque (Figure 1a) to derive a network model whose nodes are localized populations of neurons and whose edges are representative axons. Spatial parameters of the model approximately correspond to the actual two-dimensional surface view of the visual cortex, taken from [19] which in turn were derived from [18] and [34]. The model comprises 5000 nodes, each representing the neural population under a 1 mm by 1 mm piece of cortex. The nodes are distributed across a 100 mm by 50 mm model cortical sheet. Specifically, the sheet is divided into 5000 equally-sized units and a node is placed at a location chosen uniformly at random within each unit.

We represent the primary visual cortex (V1) in a 100 mm by 10 mm region of the model cortical sheet and potential secondary areas in the remaining region. The initial state of the model is shown in Figure 1b, where the location of nodes within V1 are color-coded. In the macaque cortex, the horizontally-extended blue-to-black edge represents the peripheral visual field on an unrolled cortex, and the white-to-red edge the central visual field. V1 nodes along the caudorostral axis (white-to-blue, red-to-black) span 90 degrees center-to-periphery of the visual field. The blue-to-black boundary is located at the anteriormost aspect of V1, and is curved. The representation used for this model employs the approximate ratio of the length of the peripheral border to the length of the peripheral-to-vertical meridian: essentially, the horizontal meridian is “split” and laid on the abscissa, the white-to-red axis, and span 180 degrees up-to-down of the central visual field.

We model cortical development by means of a developmental program that adds new directed edges to the network, originating at nodes in the primary visual cortex and terminating at nodes in potential secondary visual areas. The program unfolds over sequential growth steps and a constant number of edges are added to the network at each step. The source and destination nodes of a new edge are drawn from a probability distribution that is a function of two variables (1) pairwise activity correlations between nodes in the network and (2) available number of contact resources at each node. Pairwise activity correlations arise from spontaneous excitation of V1 nodes at every growth step, which in the cortex arises from multiple sources [35–37]. Excitation of a V1 node generates correlated excitation in nodes in its immediate spatial neighborhood as well as in the neighborhood of nodes where it projects edges to (Figure 1c), corresponding to a spread of neural activity outwards from excited nodes. New edges preferentially form between nodes whose activities are more correlated. As these new edges are added, the number of contact resources of the connecting nodes decline (Figure 1d), corresponding to a depletion of synaptic contact points in the respective neural populations. Model equations are provided in the *Methods* section.

Thus during the growth of the network, a new edge is more likely to form between two nodes that have higher activity correlations and more contact resources compared to other node pairs in the network. Addition of these edges alter pairwise activity correlations in the network and the available contact resources of the connecting nodes, thereby altering the probability distribution from which subsequent edges are drawn. Thus, the pairwise likelihood of new edges between nodes in the network changes in time as the network grows (Figure 1e).

## 3 Results

### Topographically organized mirror-reversing maps

The generation of maps in secondary visual areas from the “seed” map of V1 is demonstrated in Figure 2. The figure depicts the progressive formation of new topographically-organized areas at ten time points. The initial state of the network is shown at time t=0. The location of nodes in V1 are color-coded with the red-to-black and the white-to-blue progressions representing locations center-to-periphery of the visual field and the white-to-red and the blue-to-black progressions representing locations up-to-down. As the developmental program progresses in time, the visual field locations that nodes outside of V1 come to represent are also depicted by color. Specifically, each (RGB) color component of a node outside of V1 is determined by averaging the corresponding color component of its incoming edges, where each edge is assigned the color of its source node (residing in V1).

**Figure 2:**
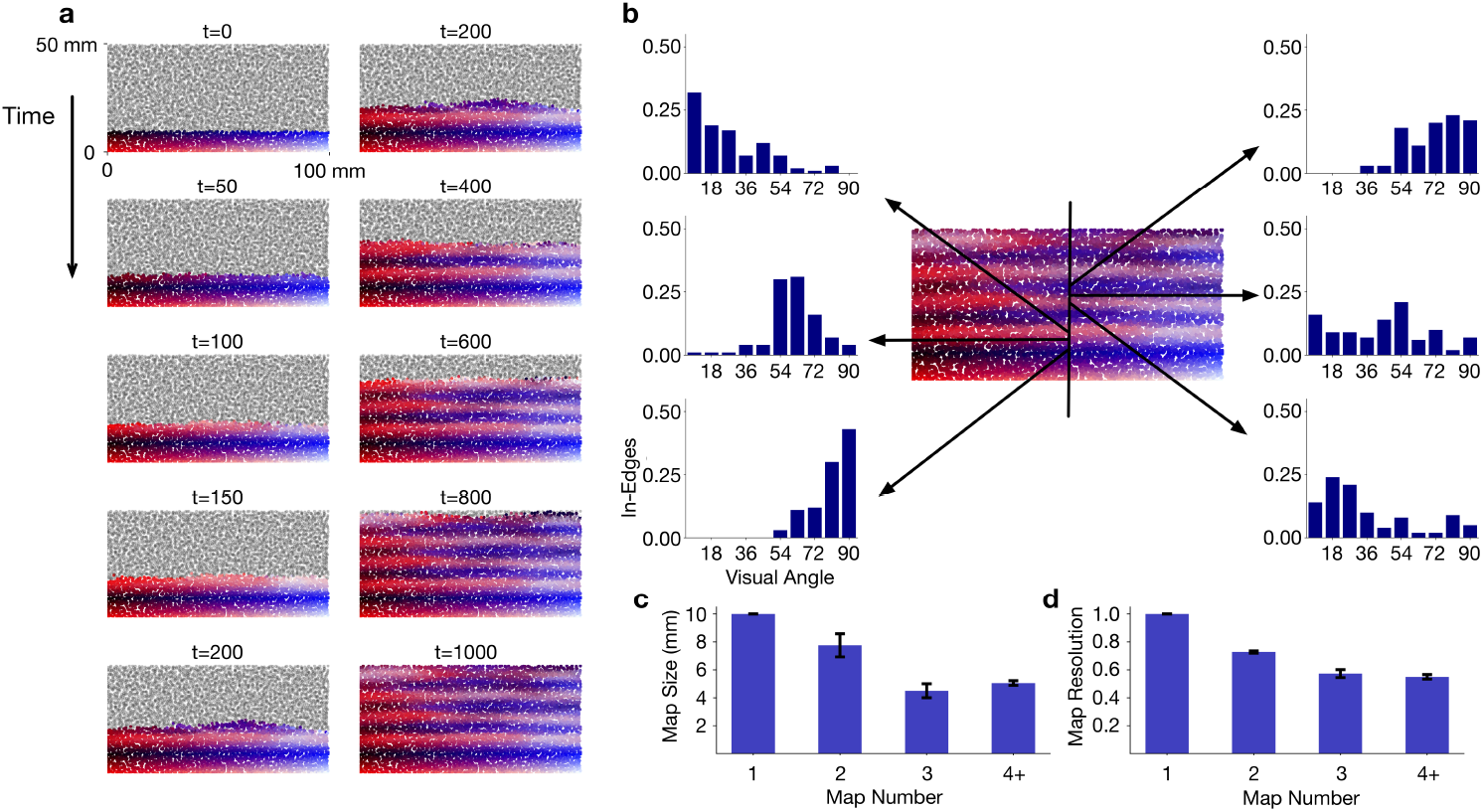
(a) Sequential steps of network growth from the initial state at time t=0 to the final state at time t=1000. At each growth step new edges are added from nodes in V1 to nodes outside of V1. The location of the center-to-periphery visual field represented by each node is color coded. The gradual change in color along the caudorostral axis depicts the progression of the topographical representation within a map and the mirror reversals across successive maps. (b) Receptive fields of six nodes along the caudorostral axis. The bar graphs depict the visual field represented at each of the nodes. Each bar shows the number of incoming edges (normalized count) from V1 nodes that represent particular center-to-periphery locations (shown as visual angles) of the visual field. (c) Size of successive maps, averaged across multiple 1 mm x 50 mm slices along the caudorostral axis. Error bars depict standard deviation. (d) Center-to-periphery receptive field resolution in successive maps. Resolution of a node is measured as 1 – σ/45, where σ is the standard deviation of the center-to-periphery visual field angles represented in the incoming edges of the node. Nodes have higher resolution when their incoming edges represent more similar points of the visual field. Values are averaged across all nodes in the maps.

The first nodes to establish synaptic contacts in “V2” are those close to V1 nodes representing peripheral locations up-to-down, because of correlated activity arising from their immediate spatial propinquity. As new edges are added, the next to establish contacts are those representing positions less peripheral, as contact resources of the most peripheral nodes decline. The mirror-reversal of the first establishment of the horizontal axis of the visual field proceeds in this fashion forward, and by t=200, the first representation of the visual field is complete, “V2”, and the next mirror-reversal emerges, this time of the vertical meridian of the central visual field, initiating “V3”. Eight mirror-reversing representations of the visual field are established by t=1000. The distribution of incoming edges at six keyed nodes at this time is shown in Figure 2B.

### Map size and resolution

Each of the successive maps arising from the growth of the network represents the full central-to-peripheral extent of the visual field. Consistent with approximate size ratios measured in the macaque, successive maps from V1 to V3 compress in size along the caudorostral axis. This is shown in Figure 2C where, on average, V2 and V3 are around 80% and 50% the size of V1 respectively. In tandem with this successive size compression, the resolution of the maps, as measured by the distribution of incoming edges in their nodes, successively falls (Figure 2D).

The successive reduction in map resolution and size arise as proximal V1 nodes with correlated activity tend to project edges to the same V2 node. Thus, each V2 node comes to represent a larger area of the visual field and have a coarser resolution compared to the point-to-point retinotopic mapping in V1; consequently, the overall V2 map is compressed relative to V1. In contrast to the high-resolution V1 nodes at the V1-V2 boundary, lower-resolution V2 nodes at the V2-V3 boundary initiates a coarse V3 map that undergoes a further reduction in resolution and size as compared to V1 and V2.

### Spatial spread of correlation envelope

As noted earlier, excitation of a V1 node in the network model is accompanied by correlated excitation in nodes of its immediate neighborhood (Figure 1c), the extent of which is defined by a two-dimensional Gaussian function parameterized by a spread along the mediolateral axis (*σ_x_*) and a spread along the caudorostral axis (*σ_y_*; *Methods*). The spatial extent of these spreads affect the degree of activity correlations between nodes, with a broader spread generating higher activity correlations compared to a narrower spread. We refer to this spread as V1 activity spread. Excitation of a V1 node also generates correlated excitation on nodes in secondary visual areas to which it projects edges to (Figure 1c). This excitation spreads to the neighborhood of the receiving node, the extent of the neighborhood being defined by a second Gaussian parameterized by spreads along each of the two axes. We refer to this as the V+ activity spread. Below we investigate map properties as the V1 and V+ activity spreads are systematically varied.

The effect of increasing the V1 activity spread while the V+ activity spread is kept fixed at an optimal value is shown in Figure 3a. The left-most map uses a Gaussian function with standard deviation *σ_x_* = 0.5 and *σ_y_* = 0.5, corresponding to a spread that falls off to 60% of its peak value within 0.5 mm along either axes. Here, eight topographic maps form from the initial V1 seed, corresponding to eight mirror reversals as depicted in Figure 3b. Note the periodic change in represented visual angle along the propagating axis. On increase to a spread of *σ_x_* = 1.0 and *σ_y_* = 1.0, the second map in the series, corresponding to a Gaussian that falls off to 60% of its peak within 1 mm, resolution of the visual map has declined, and topographic organization beyond the fifth map has essentially disappeared. A larger spread increases activity correlations between nodes in V1, resulting in V1 nodes within a spatially extended neighborhood to project to common targets, consequently reducing the resolution of visual field representations in secondary areas. When the spread increases further (fourth column in Figure 3a), notice that while the center-to-periphery periodicity of the maps virtually disappears, the up-to-down alignment of the maps remain for a few iterations.

**Figure 3:**
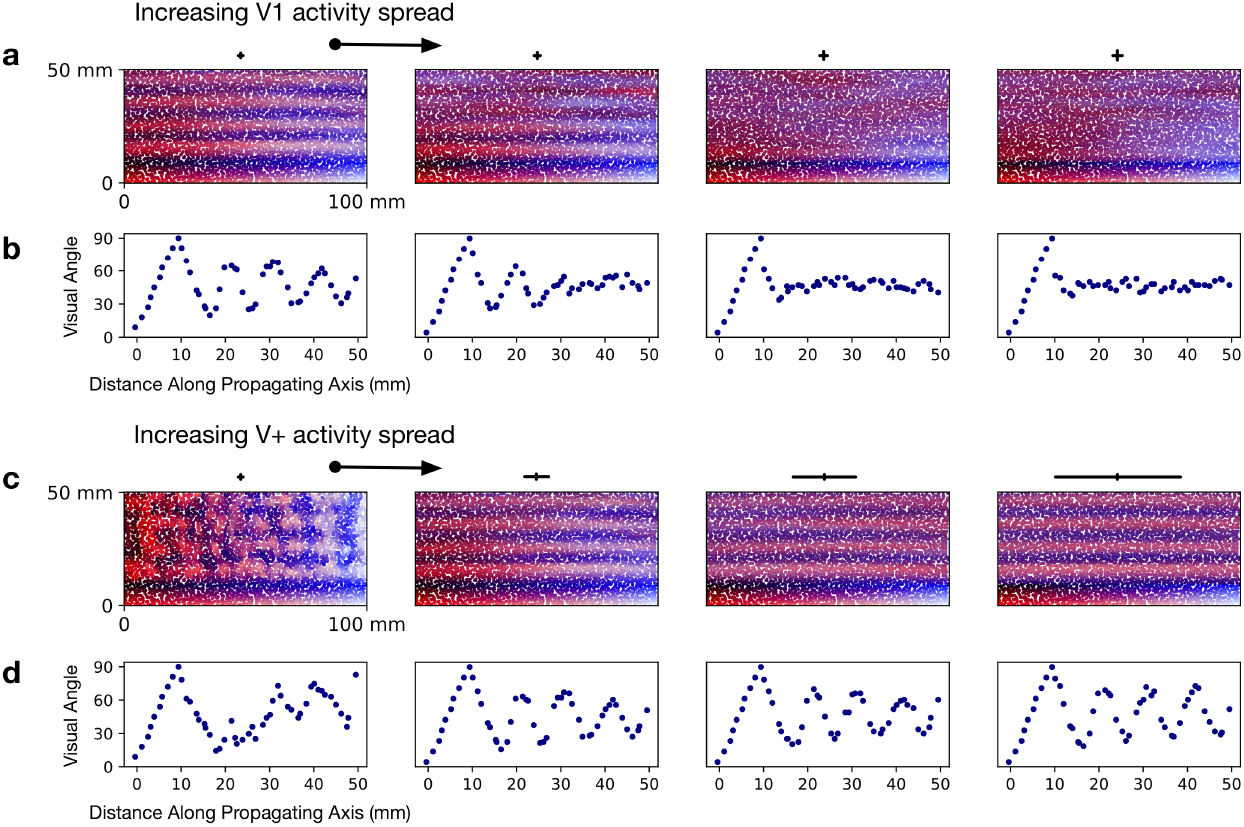
(a) Effect of increasing the activity spread in V1. Solid black lines above the color-coded maps show the spatial extent of the spread (2*σ*) along each axis. Standard deviations of the Gaussian activity spread in each of the four panels are (*σ_x_* = 0.5, *σ_y_* = 0.5), (*σ_x_* = 1.0, *σ_y_* = 1.0), (*σ_x_* = 1.5, *σ_y_* = 1.5) and (*σ_x_* = 2.0, *σ_y_* = 2.0) from left to right. The activity spread in the secondary visual areas was kept constant at an optimal value across all four panels. (b) Visual angles represented in a 1 mm x 50 mm slice along the propagating axis of each map in *a*. The visual angle represented in a node is measured as the mean of the visual angles represented in its incoming edges. (c) Effect of increasing the activity spread along the mediolateral axis in the secondary visual areas. Standard deviations of the Gaussian activity spread in each of the four panels are (*σ_x_* = 0.5, *σ_y_* = 0.5), (*σ_x_* = 5.0, *σ_y_* = 0.5), (*σ_x_* = 12.5, *σ_y_* = 0.5) and (*σ_x_* = 25.0, *σ_y_* = 0.5) from left to right. The activity spread in V1 was kept constant at an optimal value across all four panels. (d) Visual angles represented in a 1 mm x 50 mm slice along the propagating axis of each map in *b*.

The effect of increasing the V+ activity spread along the mediolateral axis while the V1 activity spread is kept fixed at an optimal value is shown in Figure 3c-d. A small spread of the Gaussian (leftmost column; *σ_x_* = 0.5 and *σ_y_* = 0.5) induces local clusters of correlations and generates disorderly maps. When the spread along the mediolateral axis is increased (second from the left column; *σ_x_* = 5.0), orderly maps emerge. This is a consequence of higher activity correlations along the mediolateral axis that establishes continuity of a particular center-to-periphery visual field location represented along this axis. Interestingly, the initial distribution of growing axons is somewhat anisotropic over the embryonic cortex [38], a potential source of these anisotropic correlations. As the spread is increased further along the mediolateral axis, map order and periodicity stays intact; thus the V1 activity spread is the dominating influence in this parameter regime.

### Scaling cortex size

As the cortex increases in size from mouse to macaque, V1 axons extend to an expanded area of the cortical surface whose parameters are described in [19]. We investigate the effects of this expansion on the properties of the maps generated by our model (Figure 4). We find that, as the cortical surface area increases, a larger number of topographically-organized secondary visual areas are generated (Figure 4a-b). The expanded area available for V1 axonal outgrowth results in repeated mirror flips of the V1 topographic map arising from the iterative mechanisms of map propagation in the developmental program. Furthermore, the increase in cortical surface area along the mediolateral axis results in reduced activity correlations between V1 nodes representing distinct up-to-down locations of the visual field. As a consequence, the likelihood of these nodes projecting to common targets is reduced, resulting in finer resolution of visual field representations in secondary visual areas of the larger cortex (Figure 3c).

**Figure 4:**
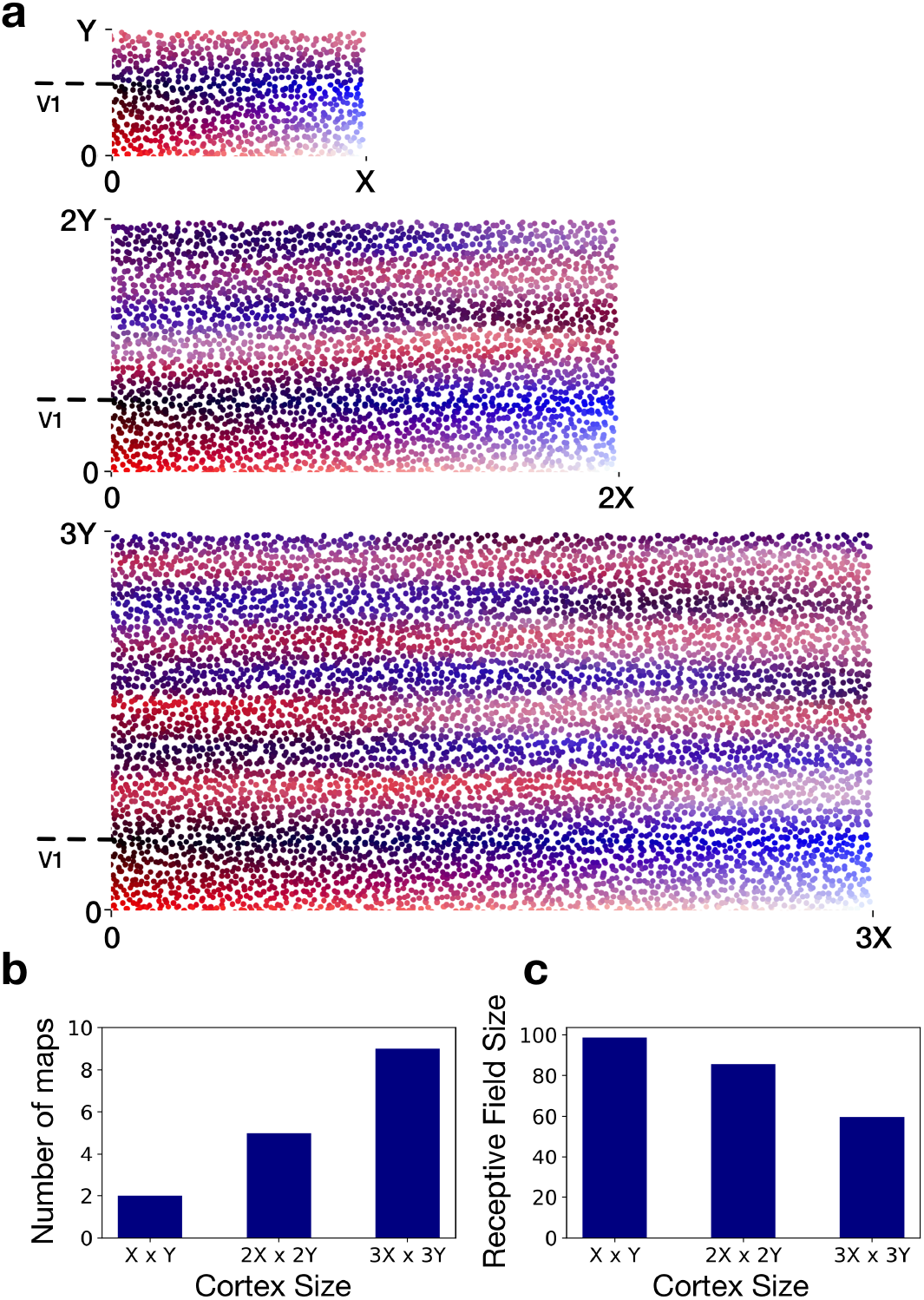
(a) As the model cortex grows in size, conserved rules of network growth generate systematic variations in cortical organization. Most notably, a larger number of topographically-organized secondary visual areas emerge as the total surface area of the cortex increases. The largest cortex shown is three-fold larger along each dimension compared to the smallest cortex. The length of V1 along the propagating axis is held constant (at 0.6*Y*) for all three cortices. (b-c) The number of maps and the average size of V2 receptive fields in the cortices shown in *a*. The receptive field size of a V2 node is measured as the standard deviation of up-to-down visual field angles represented in its incoming edges.

## 4 Discussion

### Principal findings and empirical support

The investigations presented here show that iterated topographic maps of a “seed” map can be generated from minimal information sources. The critical information source parametrically explored here is the activity correlation between neuronal populations that arise due to the formation of synaptic contacts between them.

Map development is conceptualized as extending from the seed region in successive steps, this feature to be discussed later. The first map of the secondary visual areas mirror-reverses at the border of V1 and is smaller in overall spatial dimensions than V1. Multiple mirror-reversals and maps are formed by this process, with each one smaller in spatial dimension than the prior map in the first couple of iterations, and with lesser spatial resolution at each of its nodes overall. Progressive flattening of the V1 activity spread eventually disrupted map propagation, except for some “passive” alignment of similar receptive field areas on the mediolateral axis (x-axis, Figure 3A rightmost column).

This developmental program *did not specify* any preference of edges for any particular part of the substrate (often called “axon-target interaction”) other than preferences arising from activity correlations, any recognition process between edges (often called “axon-axon recognition”) [30], nor the size of the “cortical areas” to be formed, except by the limits of the overall area of the propagating region. Receptive-fields were composed only from the convergence of the most-correlated edges.

The overall spatial parameters of the maps explored were chosen to represent the spatial parameters of the actual visual cortex in 2D form, but with no attempt to relate neuron numbers to node numbers. As mentioned, these parameters included 1) the area of the seed region, 2) the area of the region for propagation, 3) the initial spatial extent of extending axons implied by Gaussian envelopes of correlations, and 4) the large asymmetry in the length of the border over which maps propagate, compared to the shorter dimension on the propagating axis. The border of V1 with secondary visual areas corresponds to the upper-to-lower limit of the visual field on its first mirror-reversal, and the vertical meridian of the visual field on the second mirror reversal. The other axis, the “propagating” axis, at its midpoint is the horizontal meridian of the visual field, which is “cut” to lay out the map on one continuous axis on the graph, as in Figures 2–4, exactly analogous to the procedure used to lay out the curved cortical surface in Figure 1a.

In the model, the propagation of the V1 map is iterative, and proceeded along the caudorostral axis from the nodes closest to V1 to nodes most distant. This propagation was driven by activity correlations between neighboring nodes and wiring limits within nodes that had established connections. These two minimal features, activity-based attachment and wiring density limits, were essential to establish map polarity, initially forming edges between highly correlated nodes located at the V1-V2 border, and subsequently between nodes further away from the border, as contact resources of nodes at the border declined. This process propagated and ordered the rest of the V2 map and its mirror-reversing iterations. It is worth underlining that multiple additional mechanisms might contribute to map organization: a major lesson learned from multiple investigations of retinotectal map formation in multiple vertebrates, following Sperry’s initial work, was the demonstration that virtually all possible sources of order were exploited in map formation, including spatial and temporal maturational asymmetries, neuron/location recognition systems and activity dependent ordering [31].

One surprising observation of studies of both developing [38] and mature intracortical axon extent is how very large the area of overall cortex is that may be reached from a “point” origin in the cortex [18,19] (if recovered from identified single axons, the covered area becomes patchier but not larger in its outside perimeter). In the mouse, the range of projections from a point injection can reach 80-90% of the cortical surface; in the rhesus monkey, whose surface area is 200x greater than the mouse’s, the range is about 50%. The terminations of the axons from this point have an “exponential distance falloff” in which most terminations (50-90%) are close to the origin, and the remaining small fraction reach further [19]. Looking in a developing rodent, the hamster, which is born early enough so that the cortex may be injected when the final supragranular cortical neurons, the main source of intracortical projections, have only just migrated into position [26], the full range of axon extent (as a percentage of cortical surface area, which is quite small at this point) is established almost immediately [38]. There is no “front” of axon outgrowth, so the spatial correlations of activity by which the maps are found must be found within the whole axon outgrowth complement. Interestingly, the overall pattern of axon outgrowth is set up when the cortical surface is only about 20% of its adult area [26], so like most of the brain, axon stretch rather than axon extension will characterize most axon growth [39], a fact hypothesized to be of material importance in the establishment of gyri and sulci [40].

Topographically organized activation, emanating from “retinal waves” and other sources, reaches the cortex via the lateral geniculate nucleus from the moment of first thalamic innervation, and may even have an earlier influence in the transitory subplate [35]. The cortex becomes unresponsive to this input before eye opening [36], at which time the cortical areal extent has approximately doubled [26]. The emergence of the retinotopic map in V1 has been extensively studied in several species from the earliest accessible times, typically postnatal and post-eye-opening. The initially surprising, but now well-accepted result that the V1 retinotopic map is close to, or at its adult specificity at its very first emergence was quickly established (as contrasted with the stabilization of ocular dominance columns, or midbrain visuotopic maps which stabilize later). The primary sensory and sensorimotor nuclei of the thalamus and their corresponding cortical projection regions appear both temporally and informationally privileged [23, 37, 41]. These thalamic nuclei are generated before all other thalamic nuclei and establish their cortical connections first as well [26]. The neurons that will become somatosensory (or somatomotor), auditory and visual cortices are not generated prematurely, but rather are positioned at their typical relative locations within the cortical plate by rostral and caudal polarizers [42]. Once positioned, genomic studies demonstrate that the early primary sensory regions are different from all other cortical areas, replete with gene expression for surface proteins and receptors implicated in axon-target interaction and topographic map polarization and organization [7]. Both molecular and activity-dependent processes are thought to contribute to the early stabilization of the geniculocortical map.

Overall, therefore, the basic premise of this study, that primary cortex, V1, is a retinotopically-organized source of correlated activity, with axons in place across the cortical surface with the potential to organize secondary cortical areas is very well supported, in multiple species. At this point the enormous lacuna in understanding cortical development in non-primary areas presents itself. Massive research effort focused on the precise mechanisms of pre- and post-experience retinotopic organization of V1, has existed in parallel with intense interest in what the “nature” of cortical areas are. Yet, there seem to be no studies whatsoever of the early development of topographic order in extrastriate areas, and only a few demonstrating the simple presence of any secondary visual areas [43]. Notably, an early neurophysiological study, inspired by the demonstration of “face” and “hand” recognition cells in monkey inferotemporal cortex, looked for the same in infant monkeys, and found instead virtually no cells activated by visual input [44]. This null finding finds support in the very late emergence of face responsive areas anywhere in the cortex in human children in an extensive series of studies using fMRI [45].

### “Evolution” of cortical areas and organization in cortices of varying size

In general, the number of cortical areas increases with overall cortical area. In the first twenty to thirty years of cortical mapping studies, enough different species studied with rough, and roughly similar techniques could be found to estimate that the rate of increase of number of cortical areas to overall cortical area was approximately linear over the range of brain sizes represented by small rodents and shrews to carnivores and small monkeys [46, 47]. Interestingly, peripheral visual acuity interacted with this function, appearing to result in more areas. Better technical expertise, completeness and complexity in the study of a few select species have now made newer studies incomparable with older ones, so that such comparisons are no longer possible. While systematic quantitative comparisons across species are not possible, qualitative descriptions certainly are.

#### The interpretation of extrastriate regions in mice

From the outset, there has been substantial disagreement about the region of the brain surrounding primary visual cortex laterally and medially in mice (and other rodents) responsive to visual stimulation. One camp held that a number of cortical areas could be found, corresponding to the range of secondary visual areas in primates [48] while a second camp found a single V2 much like monkey, though topographically disorderly [49]. Representatives of these positions may be found to this day (specialized regions: [50]), with a new entrant, dividing the circumstriate belt into two, with its mostly medial and mostly lateral regions corresponding respectively to the dorsal and ventral streams described in primates [43]. For the purposes of this paper, the only necessary features of mouse extrastriate cortex are a small region of visually-responsive cortex abutting the V1 border, of uncertain topographic organization.

#### Characteristic changes in organization in larger brains

The pattern of change in organization of areas in larger brains is drawn mostly from the comparison of rhesus macaques, humans, and several New World (South American) monkeys [4, 32, 51]. The number of cortical areas increases, and generally each area contains a representation of the entire visual field (with a few debated cases). For the first several areas, a clear mirror-symmetric replication is observed (V1, Center to Periphery; V2, Periphery to Center; V3, Center to Periphery), each map smaller in surface area than the preceding one. Thereafter, the topographic ordering may have become so degenerate (for example, in some lateral parietal areas, every receptive field may represent the center of gaze) that mirroring may not be possible to detect. Nevertheless, cortical areas may be organized in rough hierarchy by virtue of their pattern of feedback versus feed-forward circuitry, and corresponding architectonics, showing that decrease in overall size continues to an asymptote. How are we to understand this addition of cortical areas?

### Conserved rules of development inform the interpretation of new cortical areas

If the model described here proves correct, it can constrain the interpretation of what a “new” area means. Proof is required: it will be absolutely essential to demonstrate the nature of development of secondary cortical areas in an animal with a large enough brain to produce several orderly retinotopic maps, and show that it is correlated activity, and not molecular pre-specification of connectivity (as seen in V1) that produces them. New cortical areas would thus be the outcome of the changing geometry of regularly scaling brains. In a small extrastriate area, only one orderly map can be supported, mirror-flipped, V2; in a slightly larger one, two maps, V2 and V3, and so forth. The question of homology of areas across species can be at least partially resolved. In a larger brain with an additional visual cortical area, it cannot be said that a “new cortical area” has been specified. Rather, a larger extrastriate region has been produced, by allometrically predictable enlargement of the cortex, and that region has been subdivided by activity-dependent self-organization into three, rather than two retinotopic maps, or five instead of three, and so forth. The most distal map from V1 is not the “new” area, as all areas have undergone reorganization, it is simply the most distal division of the whole reorganized region. Nevertheless, it is not hard to see how such an underlying process, in the context of overall hierarchical organization of the cortex, might serve as a mechanism by which a new regularity in the pattern of sensory input to the whole organism, or a particular pattern of experience could produce new computational possibilities.

## 5 Methods

### Physical composition of the model

The visual cortex of the rhesus macaque is modeled as a network comprising 5000 nodes, each representing localized neural populations in a 1 mm by 1 mm piece of cortex. The nodes are distributed across a 100 mm by 50 mm model cortical sheet. Specifically, the sheet is divided into 5000 equally-sized units and a node is placed at a location chosen uniformly at random within each unit.

The primary visual cortex is represented in a 100 mm by 10 mm region of the model cortical sheet. The network is programmed to grow in sequential steps by adding new directed edges from nodes in the primary visual cortex to nodes within the rest of the cortical sheet. This represents the formation of new synaptic contact points between neurons in the respective populations as the cortex develops. The relative dimensions of primary and secondary cortical regions and their borders conform approximately to those observed for the rhesus macaque cortex (Figure 1a-b).

### Parameters determining activity correlation

At every growth step, each of the nodes of the model V1 spontaneously generate a unit level of excitation (equal to 1), corresponding to spontaneous neural activity during development. These unit excitations are generated one node at a time and their specific order is random. Unit excitation in one V1 node is accompanied by correlated excitation in neighboring V1 nodes (Figure 1c). Specifically, correlated excitation *a_j_* on V1 node *j* arising from a unit level of excitation in V1 node *i*, is determined by a two-dimensional Gaussian function:

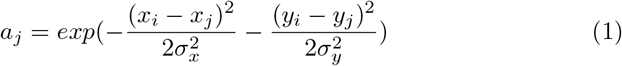

Here, *x_i_* and *x_j_* are the respective positions of nodes *i* and *j* along the mediolateral axis of the cortical sheet, *y_i_* and *y_j_* are their respective positions along the caudorostral axis, and *σ_x_* and *σ_y_* are parameters that determine the spread of the Gaussian along each of the axes.

Excitation of V1 nodes also generate correlated excitation in nodes within the rest of the model cortical sheet. Specifically, unit excitation of a V1 node *i* generates excitation on node *j* that it projects edges to (Figure 1c) in proportion to the the number of projected edges. These edges also induce excitation on nodes residing in the neighborhood of *j*, based on a Gaussian function of distance from *j*. This function takes the same form as the right-hand side of Equation 1, but is parameterized independently. It represents a spread of excitation outwards from node *j*.

Thus, unit excitation of a V1 node results in pairwise activity correlations between all nodes in the network. Given unit excitation of a V1 node, the activity correlation between two nodes *i* and *j* is computed as *a_i_a_j_*, where *a_i_* and *a_j_* are resulting excitation levels of node *i* and node *j* respectively. In a given growth step, the net activity correlation *c_ij_* between nodes *i* and *j* equals the sum of their activity correlations as unit excitations are generated in each of the V1 nodes, one at a time.

### Network Growth

Based on these activity correlations, the network generates new edges emanating from nodes within V1 and terminating on nodes that reside outside of V1. New edges are drawn from a probability distribution that evolves as the network grows. Specifically, at each growth step, the probability *p_ij_* of a new edge from node *i* to node *j* equals

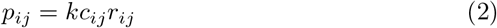

where *c_ij_* is the net activity correlation between node *i* and node *j*, *r_ij_* is a measure of the available contact (synaptic) resources in the two nodes, and *k* is a normalization constant. Thus new edges preferentially form between nodes that have higher activity correlation and more available contact resources.

The measure of available contact resources *r_ij_* between nodes *i* and *j* in Equation 2 is defined as *r_ij_* = *a_i_d_j_*, where *a_i_* is a measure of available axonal contact resources in node *i* and *d_j_* is a measure of available dendritic contact resources in node *j*. Both these terms follow a logistic decay as edges are added to nodes *i* and *j* (Figure 1d). Specifically,

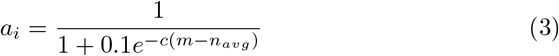

where *m* is the number of outgoing edges of node *i*, *n_avg_* is the number of outgoing edges of V1 nodes on average (which increases as the network grows), and c is a constant set to 0.1. Similarly,

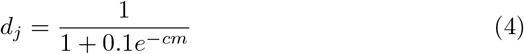

where *m* is the number of incoming edges of node *j* and *c* is a constant set to 0.05.

The network is initialized without edges and grows in sequential growth steps, with a constant number of edges added in each step. The source and destination nodes of each of these edges are independently drawn from the probability distribution of Equation 2. The probability distribution evolves as the network grows (Figure 1e), as new edges alter pairwise activity correlations and the available contact resources in the network.

This developmental program of the early visual cortex, in addition to serving as our growth model in the present study, can also serve to self-organize a new generation of parallel computing systems that scale as cortexlike sheets of arbitrary size [52–54].

